# Sperm activation for fertilization requires robust activity of the TAT-5 lipid flippase

**DOI:** 10.1101/2025.03.06.641851

**Authors:** Katherine A. Maniates, Saai Suryanarayanan, Alissa Rumin, Morgan Lewin, Andrew Singson, Ann M. Wehman

## Abstract

During fertilization, sperm and egg membranes signal and fuse to form a zygote and begin embryonic development. Here, we investigated the role of lipid asymmetry in gametogenesis, fertilization, and embryogenesis. We find that phosphatidylethanolamine asymmetry is lost during meiosis prior to phosphatidylserine exposure. We show that TAT-5, the P4-ATPase that maintains phosphatidylethanolamine asymmetry, is required for both oocyte formation and sperm activation, albeit at different levels of flippase activity. Loss of TAT-5 significantly decreases fertility in both males and hermaphrodites and decreases sperm activation. TAT-5 localizes to the plasma membrane of primary spermatocytes but is sorted away from maturing spermatids during meiosis. Our findings demonstrate that phosphatidylethanolamine asymmetry plays key roles during gametogenesis and sperm activation, expanding the roles of lipid dynamics in developmental cell fusion.

## Introductio

Fertilization initiates embryonic development and depends on the formation of functional gametes. Oocytes and sperm must recognize each other and fuse their membranes to form the zygote (Deneke and Pauli, 2021; Krauchunas et al., 2016). In mammals, phosphatidylserine (PS) lipids found in the cytofacial leaflet of the plasma membrane are exposed to the surface of sperm and PS-binding receptors are required in the egg plasma membrane for fertilization (Rival et al., 2019), indicating that PS exposure and recognition promote sperm-egg fusion. The regulated asymmetry and symmetry of lipids across membrane bilayers is necessary for many processes, including vesicle release, signaling, cell fusion, and cell corpse clearance (Andersen et al., 2016; Folmer et al., 2009). However, PS lipids are only one of several classes of lipids normally enriched in the cytofacial leaflet and exposed during cell fusion (Lorent et al., 2020). Phosphatidylethanolamine (PE) lipids have also been implicated in cell fusion, and PE is exposed on capacitated sperm (Irie et al., 2017; Vries et al., 2003), but a function for PE exposure on sperm has not been studied.

The asymmetric distribution of lipids is facilitated by flippase proteins that enrich specific lipids in the cytoplasmic leaflet of the plasma membrane. The P4-ATPase family of proteins hydrolyze ATP to flip the hydrophilic headgroup of phospholipids across the hydrophobic barrier of the membrane (Norris et al., 2024). To examine the role of PE lipid asymmetry during spermatogenesis and fertilization, we focused on the function of the essential P4-ATPase, TAT-5. The ATPase activity of TAT-5 is required to maintain the asymmetry of phosphatidylethanolamine (PE) on cell surfaces, and loss of *tat-5* results in sterility and embryonic lethality in *C. elegans* hermaphrodites (Wehman et al., 2011). TAT-5 is conserved from yeast to mammals, and mouse knockouts for the TAT-5 ortholog ATP9A are both male and female infertile (Meng et al., 2023a). Thus, it is important to determine what roles TAT-5 and PE asymmetry have in male and female germ cells.

During spermatogenesis in animals including *C. elegans,* mouse, and humans, sperm shed organelles and remodel their cytoplasm to become efficient for fertilization (Breucker et al., 1985). Secondary spermatocytes undergo incomplete cytokinesis and traffic specific organelles and cytosolic proteins, including ribosomes, actin, and tubulin, to a growing structure called the residual body (Hu et al., 2019). Following meiosis II, spermatids bud off from the residual body, leaving the spermatids with the nucleus, mitochondria, Major Sperm Proteins, and Fibrous Body-Membranous Organelles (Ellis and Stanfield, 2014). The detached spermatids then undergo post meiotic sperm differentiation, also known as spermiogenesis or sperm activation. This is where sperm go from immotile spermatids to active spermatozoa with a pseudopod or flagella depending on the organism (Nelson et al., 1982; Nelson and Ward, 1980). After spermatid detachment, PS is exposed on the surface of the residual body which triggers residual body removal by phagocytic pathways that mediate apoptotic cell removal (Huang et al., 2012). However, PE dynamics during spermatogenesis and sperm activation have not yet been examined.

In this work, we find that TAT-5 is required for both male and female fertility in *C. elegans.* We show that TAT-5 localizes to the plasma membrane of primary and secondary spermatocytes and is trafficked to the membrane of the residual body. We also find that PE is exposed on the surface of the residual body prior to spermatid budding, while PS exposure occurs after spermatids separate. Both *tat-5* null and partial loss of function mutant males show reduced fertility and reduced sperm activation. Even a partial loss of TAT-5 activity caused improper localization of PE on the sperm surface. This work describes a new role of a P4-ATPase during spermiogenesis.

## Materials and Methods

### *C. elegans* strains and culture conditions

Strains were cultured on *C. elegans* growth media MYOB plates with OP50 bacteria as modified from (Brenner, 1974). Analysis was conducted at 20°C for all fertility experiments, except *fem-1(hc17ts)* which are temperature sensitive, and the culture conditions are described in the sperm migration section. Fluorescent reporter strains were cultured on NGM plates with OP50 bacteria at 23°C for imaging. Any strains containing *tat-5(tm1741)* or *tat-5(D244T)* were balanced with the *tmC18[dpy-5(tmIs1200)]* balancer chromosome (Dejima et al., 2018). Homozygous mutant animals were selected by picking animals without *myo-2::gfp* expression in the pharynx. For the sperm migration experiment, adult *fem-1(hc17ts)* animals were transferred to a new plate and shifted from 16°C to 25°C where they laid embryos. Once the resulting embryos reached the young adult stage, they were used in the experiment. A list of all strains used can be found in Supplemental Table 1.

### Genotyping

All *C. elegans* sequences were from Wormbase (Sternberg et al., 2024). Genotyping was done as previously described (Maniates et al., 2023) using the primer sequences listed in Supplemental Table 2.

### Phenotypic analysis

#### Brood size

One homozygous L4 hermaphrodite animal was placed on an individual plate to lay progeny. This animal was transferred to a new plate every 24 hours for its reproductive lifetime. All live progeny, unhatched embryos, and unfertilized oocytes were counted.

#### Hermaphrodite fertility analysis after mating with male sperm

To determine if reduced hermaphrodite fertility was due to defects solely in the sperm or also due to oocyte or embryonic lethality defects, one L4 hermaphrodite was crossed with four *cylc- 2::mNeonGreen(mon2); him-5(e1490)* males. After 48 hours, it was assessed if sperm was transferred by the presence of mNeonGreen fluorescence in the hermaphrodite. For any plates where the male successfully transferred sperm, the presence of cross progeny was scored four days after setting up the cross. To quantify the extent that hermaphrodite fertility contributed to the observed phenotypes, we then quantified the number of live larval progeny one animal was able to generate after mating with four *cylc-2::mNeonGreen(mon2); him-5(e1490)* males.

#### Male fertility

One control or mutant male and one *dpy-11(e224)* hermaphrodite were placed together on a plate and allowed to mate for 24 hours. After 24 hours, the hermaphrodite was transferred to a new plate every 24 hours until it stopped making any progeny. The number of Dpy and non-Dpy progeny was scored to determine male fertility rates.

#### Sperm activation

L4 males were placed on plates without any hermaphrodites for 24 hours. The males were dissected on a Histobond slide (VWR, 16005-108) in either Sperm Media (Singaravelu et al., 2011) or Sperm Media plus 200 μg/ml pronase. A coverslip was placed, and the slide was incubated for 5-10 minutes. Sperm were then observed using DIC.

#### Sperm migration

To assess the ability of male sperm to locate and crawl towards the spermatheca, one L4 male and one young adult *fem-1(hc17ts)* hermaphrodite were placed together on a mating plate and allowed to mate overnight at 20°C. The next day, the hermaphrodite animal was stained with DAPI (described below), and the location of the sperm was assessed.

#### Off center nuclei

To assess the off centered nuclei phenotype, male sperm dissected in sperm media were examined as described in (Shakes and Ward, 1989a).

### DAPI staining

DAPI staining was completed as previously described (Maniates et al., 2023) where animals were fixed with methanol and then stained with Vectashield Plus DAPI (H-1200-10, Vector Laboratories).

### Light and fluorescence microscopy

Images of live animals were collected on a Zeiss Axio Observer 7 microscope with a Plan-Apo 20X 0.5 NA oil objective and Excelitas Technologies X-Cite 120LED Boost illumination using a Hamamatsu ORCA-Fusion sCMOS camera controlled by 3i SlideBook6 software. Images for DAPI stained animals were collected on a Zeiss Universal microscope using a 20x objective with 0.75 NA with a ProgRes camera (Jenoptik) using ProgRes CapturePro software. Sperm activation images were collected on the Universal microscope using a 40x objective with 0.75 NA

### Localization of GFP::TAT-5 during spermatogenesis

Fluorescence microscopy for the *gfp::tat-5(wur36); him-5(e1490)* strain and corresponding *him-5(e1490)* controls were completed using a Zeiss Elyra7 Lattice Structured Illumination Microscope (SIM2) using a 488nm laser and a 63x water objective with a 1.2 NA.

### Lipid staining and quantification

Unmated adult day 1 or 2 males were dissected in 4-well slides (ibidi) in sperm media. Live sperm were stained for 30 minutes in 1:200 Alexa488-AnnexinV (Invitrogen), 0.1 µM Duramycin-LC-Biotin (MTTI), 1.5 µg/mL Alexa594-Streptavidin (Invitrogen) in sperm media. Stained sperm were imaged in sperm media on a Zeiss Axio Observer 7 microscope with a Plan-Apo 40X 1.4 NA oil objective and Excelitas Technologies X-Cite 120LED Boost illumination. Images were collected with a Hamamatsu ORCA-Fusion sCMOS camera controlled by 3i SlideBook6 software. Mean fluorescence intensity of a 3-pixel line was measured around the surface of spermatids using FIJI (NIH) and a neighboring region not containing cells. In addition, a circle smaller than the spermatid (typically ∼5 µm radius) was used to measure the mean fluorescence intensity of the cytoplasm. The neighboring background fluorescence was subtracted from the spermatid surface fluorescence or the cytoplasm fluorescence.

## Results

### TAT-5 is required for fertility in hermaphrodites

As the ATPase activity of TAT-5 is required to produce embryos (Wehman et al., 2011), we wanted to determine how TAT-5 ATPase activity impacts fertility. A large-scale screen found that *tat-5* knockdown resulted in visible defects in hermaphrodite gonads (Green et al., 2011), so we crossed a germ line plasma membrane reporter into two strains with point mutations in the conserved DGET motif of the Actuator domain of P4-ATPases: a partial loss-of-function mutant, *tat-5(D244T)*, and an ATPase-dead mutant, *tat-5(E246Q)* (Figure 1A, (Pitts et al., 2023). In control and *tat-5(D244T)* mutant hermaphrodites, embryos are visible in the uterus by DIC (Figure 1B-C), while most *tat-5(E246Q)* mutants lacked embryos (Figure 1D). In all strains, mCherry::PH(PLC1∂1)::CTPD(OMA-1) localized to the plasma membrane in both the distal and proximal germ cells, including the large cylindrical oocytes (Figure 1). The oocytes appeared normal in *tat-5(D244T*) mutants (Figure 1C), but the regular sizing and spacing of oocytes was disrupted in *tat-5(E246Q)* mutants (Figure 1D-E). To characterize the cells in the uterus of rare *tat-5(E246Q)* mutants with potential embryos, we examined whether the cells expressed mCh::PH::CTPD. The CTPD degron causes degradation of the mCh::PH::CTPD membrane reporter in fertilized embryos after the 1-cell stage (Beer et al., 2019). Unfertilized oocytes in the uterus of *tat-5(E246Q)* mutants continue to express mCh::PH::CTPD, while the fertilized embryos appear as dark ovals (Figure 1E). The presence of unfertilized oocytes in *tat-5(E246Q)* mutants suggested that TAT-5 and PE asymmetry may play a role during fertilization in addition to oogenesis.

**Figure 1.**
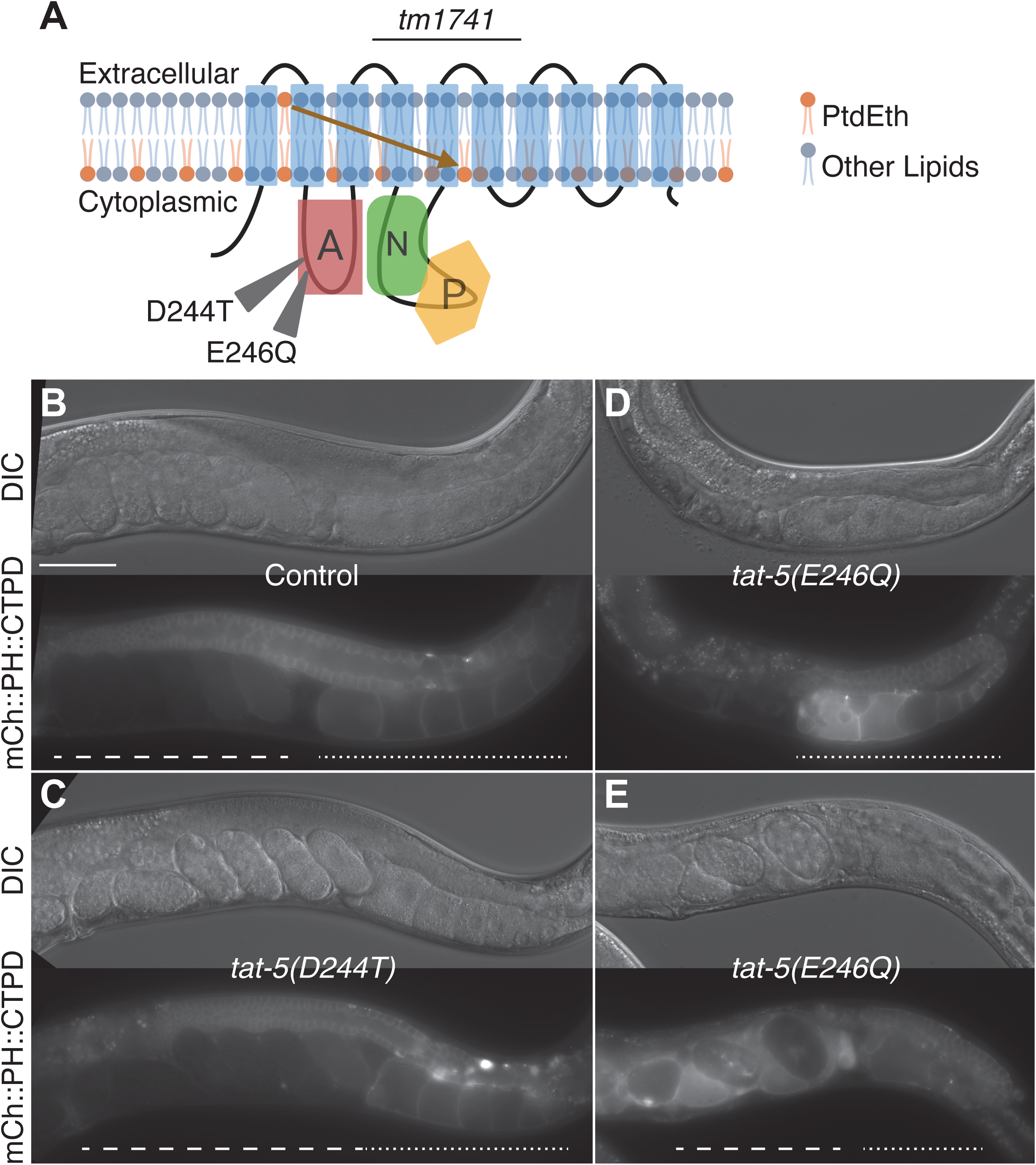
Hypomorphic and null alleles of *tat-5* have different effects on oogenesis and fertilization. A. Schematic of the TAT-5 protein adapted from (Mark et al., 2013). A-actuator domain, N-nucleotide binding domain, P-phosphorylation domain. Orange lipids indicate PtdEth lipids. Arrow denotes the flippase activity. Arrowheads indicate point mutations in DGET motif and the *tm1741* line indicates the deleted region. B-E. Expression of mCherry::PH::CTPD in control (B), *tat-5(D244T)* (C), and *tat-5(E246Q)* (D-E) young adult hermaphrodites. The length of the uterus containing embryos is underlined with a dashed line to the vulva, while the proximal germ line containing oocytes is underlined with a dotted line. Scale bar: 50μm.

### Differential requirements of TAT-5 activity for sperm and embryonic development

To ascertain the full effect that TAT-5 has on fertility, we counted the brood size of a strain with a deletion that acts as a null mutation, *tat-5(tm1741)* (Wehman et al., 2011). The *tat-5* null mutants produced no live larval progeny (Figure 2A), in contrast to wild type animals (N2), which produced on average 273 larval progeny (Figure 2A). Instead, the average *tat-5* null mutant produced 7 unhatched embryos and 29 unfertilized oocytes (Figure 2B-C), confirming that there are defects in germ cell development in addition to embryogenesis. The number of unhatched embryos and unfertilized oocytes was not significantly different than wild type (Figure 2B-C) but combined with the absence of larval progeny (Figure 2A), these data show that the germ line in *tat-5* null mutants produces significantly fewer ovulation events (Figure 2D).

**Figure 2.**
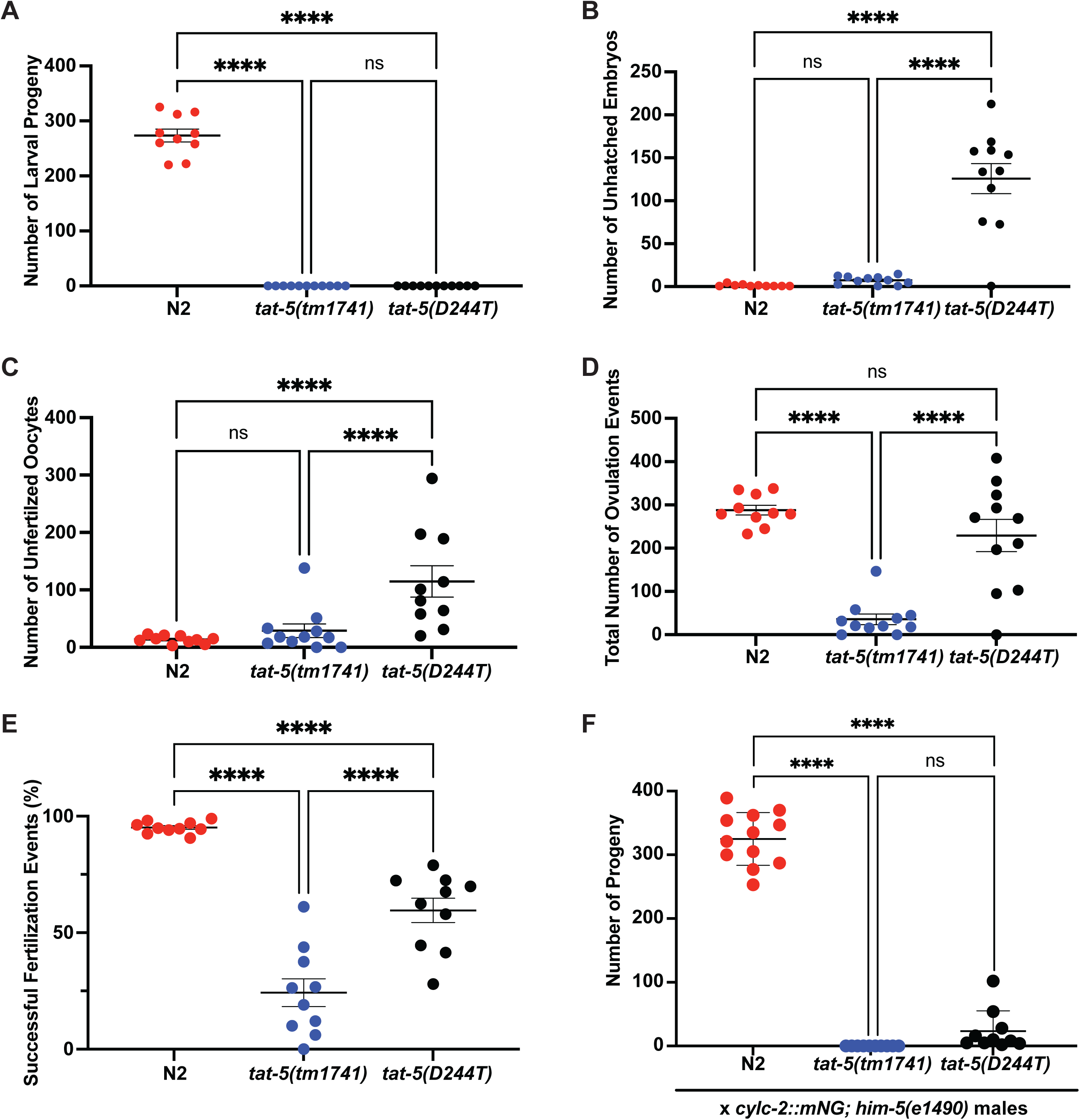
TAT-5 is required for female fertility and embryonic development. A-E. Progeny analysis of 10 N2, 11 *tat-5(tm1741),* and 11 *tat-5(D244T)* hermaphrodites. Each point represents one animal analyzed. A. Number of live larval progeny. B. Number of unhatched embryos. C. Number of unfertilized oocytes. D. Total number of ovulation events corresponds to the sum of larval progeny, unhatched embryos, and unfertilized oocytes (A-C). E. Percentage of successful fertilization events (live larval progeny and unhatched embryos compared to total number of ovulation events). F. Number of live larval progeny produced after mating with *cylc-2::mNG; him-5(e1490)* males. n=12 for N2 and n=14 for *tat-5(tm1741),* and *tat-5(D244T)*. Error bars ± SEM. One way ANOVA, **** p<0.0001, ns p>0.05.

We then analyzed the brood size of the *tat-5(D244T*) partial loss of function allele, which is predicted to lead to a 3-fold loss in ATPase activity and lipid flipping (Coleman et al., 2012). The *tat-5(D244T*) mutant hermaphrodites did not produce live larval progeny, similar to the null allele (Figure 2A). However, *tat-5(D244T*) mutants retained significant fertility, averaging 125 unhatched embryos and 114 unfertilized oocytes, significantly more than wild type or the null allele (Figure 2B-C). Comparing the total number of oocytes that passed through the spermatheca, *tat-5(D244T*) mutants did not have significantly reduced ovulation events (Figure 2D). These data indicate that a low level of TAT-5 activity is sufficient for oocyte development but not for embryogenesis.

Both *tat-5(D244T*) and null mutants showed a significant decrease in the percentage of successful fertilization events (Figure 2E), suggesting that TAT-5 also plays a role in fertilization. To understand the differential contribution of TAT-5 in sperm function and embryonic development, we mated hermaphrodites with *cylc-2::mNG*; *him-5* males for two days to determine whether male sperm could recover fertility in *tat-5* mutants. Wild type N2 hermaphrodites were able to produce cross progeny that was sired by male sperm 100% of the time (n=10), as expected since *C. elegans* preferentially use male sperm when available(LaMunyon and Ward, 1995; Singson et al., 1999). Only one of the mated *tat-5* null mutant hermaphrodites was able to generate any larval cross progeny (7%, n=14), suggesting that defects in the hermaphrodites are the primary contributor to the decreased brood size. When *tat-5(D244T)* mutant hermaphrodites were mated (n=10), 90% were able to generate some cross progeny, again showing that *tat-5(D244T)* mutant hermaphrodites produce functional oocytes.

Given how cell morphology is disrupted in *tat-5* mutants before the embryo starts zygotic transcription (Wehman et al., 2011), we wanted to determine the extent that fertilization by male sperm with wild type TAT-5 could recover embryogenesis. We set up crosses with *cylc-2::mNG*; *him-5* males and quantified the number of fluorescent larval progeny from mated hermaphrodites. We did not observe any live progeny from ten mated *tat-5* null mutants (Figure 2F). This is consistent with our previously observed defects in oogenesis and suggest that the sterility in *tat-5* null mutants is primarily due to oocyte defects.

We also examined the progeny numbers resulting from mating *cylc-2::mNG*; *him-5* males with *tat-5* partial loss-of-function mutant hermaphrodites. Although most mated *tat-5(D244T)* mutant hermaphrodites had larval progeny (Figure 2F), the number of live progeny from mated *tat-5(D244T)* mutant hermaphrodites was significantly decreased compared to wild type (Figure 2F). Considering the hundreds of ovulation events in *tat-5(D244T)* hermaphrodites (Figure 2D), these dozens of larval progeny indicate that zygotic *tat-5(wt)* is rarely able to rescue embryos from maternal *tat-5(D244T)* mutants. However, mating *tat-5(D244T)* hermaphrodites does increase the number of live progeny from zero (Figure 2A) to an average of 23 larvae (Figure 2F). Therefore, there are a fraction of cases where zygotic *tat-5(wt)* can be sufficient to rescue embryogenesis in a partial loss-of-function background. These results suggest that TAT-5 ATPase activity has differential requirements in sperm development and embryogenesis.

### TAT-5 localization is dynamic in the male germ line

To better understand the roles of TAT-5 in germ cell development and fertilization, we examined where TAT-5 localizes in hermaphrodite and male germ lines. As operonic expression is common in the hermaphrodite germ line (Reinke and Cutter, 2009), we knocked GFP into the N-terminus of the *tat-5b* and *tat-5d* isoforms (Park et al., 2024), hereafter GFP::TAT-5b, which is predicted to be expressed from its own promoter as well as from an operon (Sternberg et al., 2024). We observed that GFP::TAT-5b appeared brighter in the hermaphrodite spermatheca than in neighboring oocytes (Figure 3A), suggesting that TAT-5 could be enriched in sperm or the spermatheca. To distinguish between these possibilities, we crossed the GFP knock-in to *him-5* mutants to generate males expressing GFP::TAT-5b. GFP::TAT-5b fluorescence was first observed in the adult male germline on the plasma membrane of the meiotic region of the germ line (Figure 3B) and appeared to disappear after the division zone (arrowhead in Figure 3B), suggesting that TAT-5b in the hermaphrodite is more likely to be enriched in the spermatheca than in mature spermatids.

**Figure 3.**
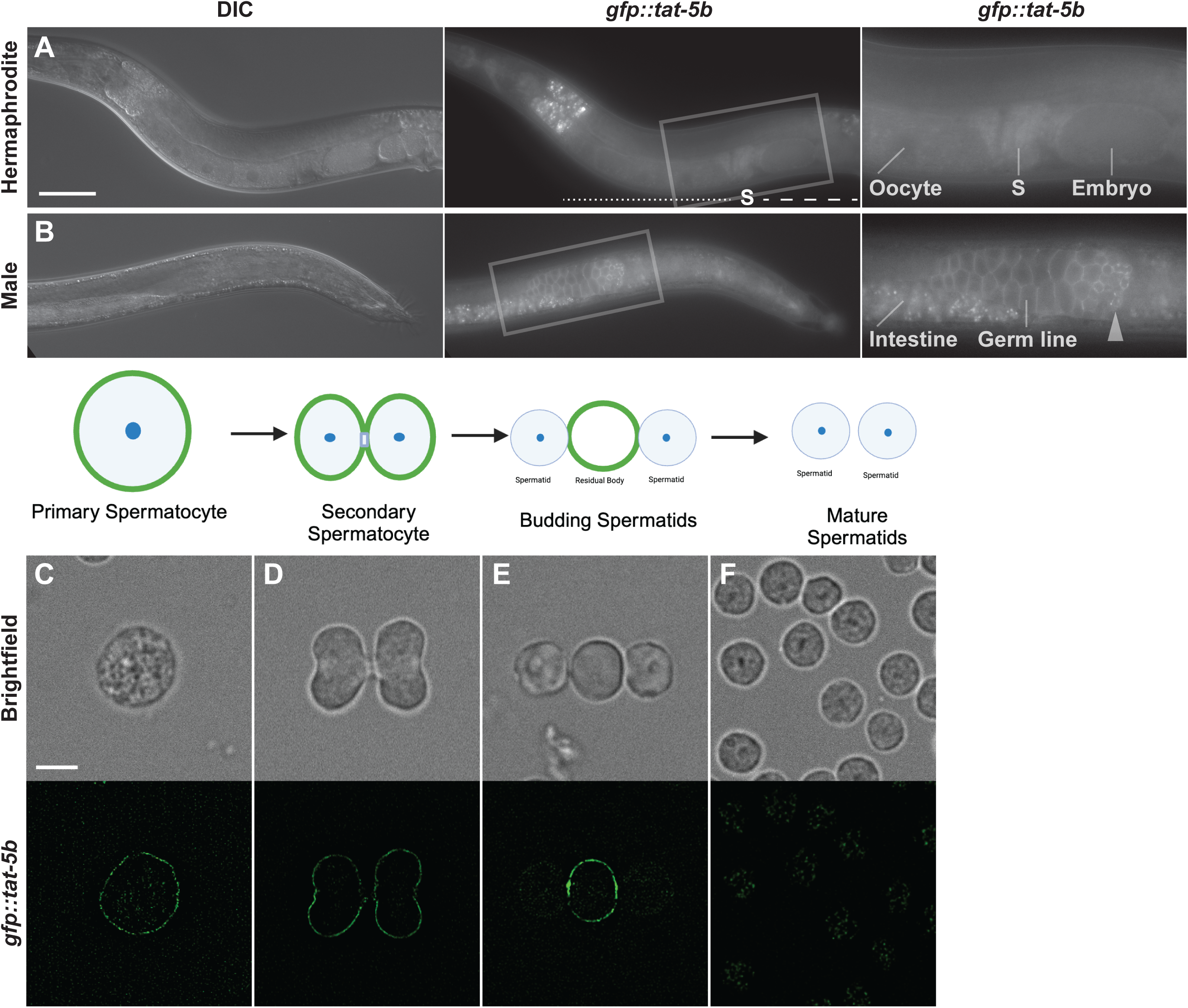
TAT-5 is expressed in both male and hermaphrodite germlines. A. GFP::TAT-5b expression in young adult hermaphrodites, including the spermatheca (S). The uterus containing embryos is underlined with a dashed line to the vulva, while the proximal germ line containing oocytes is underlined with a dotted line. Scale bar represents 50 μm for left and center panels, 25 µm for right panel. B. GFP::TAT-5b expression in young adult *him-5* males. The arrowhead indicates the division zone where spermatocytes complete meiosis. C-F. Expression of GFP::tat-5b in spermatogenic cells: C. primary spermatocyte, D. secondary spermatocyte, E. residual body, F. spermatid. Scale bar represents 5 μm.

To determine where TAT-5 localized through different stages of spermatogenesis, we dissected GFP::TAT-5b-expressing males and imaged their sperm. We found that GFP::TAT-5b localized to the plasma membrane in primary and secondary spermatocytes (Figure 3C-D), consistent with the bright fluorescence in the meiotic region of the male germ line (Figure 3B). However, GFP::TAT-5b was later enriched in the residual body membrane in comparison to the membrane of budding spermatids (Figure 3E). Consequently, GFP::TAT-5b was no longer visible on the plasma membrane of mature spermatids (Figure 3F), appearing similar to mature spermatids that do not express GFP (Supplemental Figure 1). These data are consistent with the absence of GFP::TAT-5b fluorescence in the proximal male gonad where mature spermatids reside (right of the arrowhead in Figure 3B). The trafficking of TAT-5 from the sperm membrane to the residual body suggests that TAT-5 is likely to act early during spermatogenesis.

### TAT-5 activity prevents PE exposure on the surface of spermatids

Given the role of TAT-5 in maintaining phosphatidylethanolamine (PE) asymmetry on the plasma membrane of embryonic blastomeres and the distal germ line of hermaphrodites (Wehman et al., 2011), we wanted to determine whether a decrease in TAT-5 ATPase activity also causes PE exposure on sperm. To examine PE exposure, we stained dissected sperm with duramycin, a non-permeable PE-binding lantibiotic (Navarro et al., 1985). Duramycin staining appeared as puncta on the surface of *him-5* control spermatids (Figure 4A), consistent with mature spermatids maintaining PE asymmetry on most of their plasma membrane. In contrast, we observed a >5-fold increase in duramycin staining on spermatids dissected from *tat-5(D244T); him-5* males (Figure 4B-C), with the staining appearing smooth along the cell surface of the partial loss-of-function mutants (Figure 4B). These results imply that high levels of TAT-5 ATPase activity are required to maintain lipid asymmetry in mature spermatids. As GFP::TAT-5b was not observed on the plasma membrane of mature spermatids (Figure 3E-G), this suggests that TAT-5 is needed to establish PE asymmetry in budding sperm and that mature spermatids inherit PE asymmetry.

**Figure 4.**
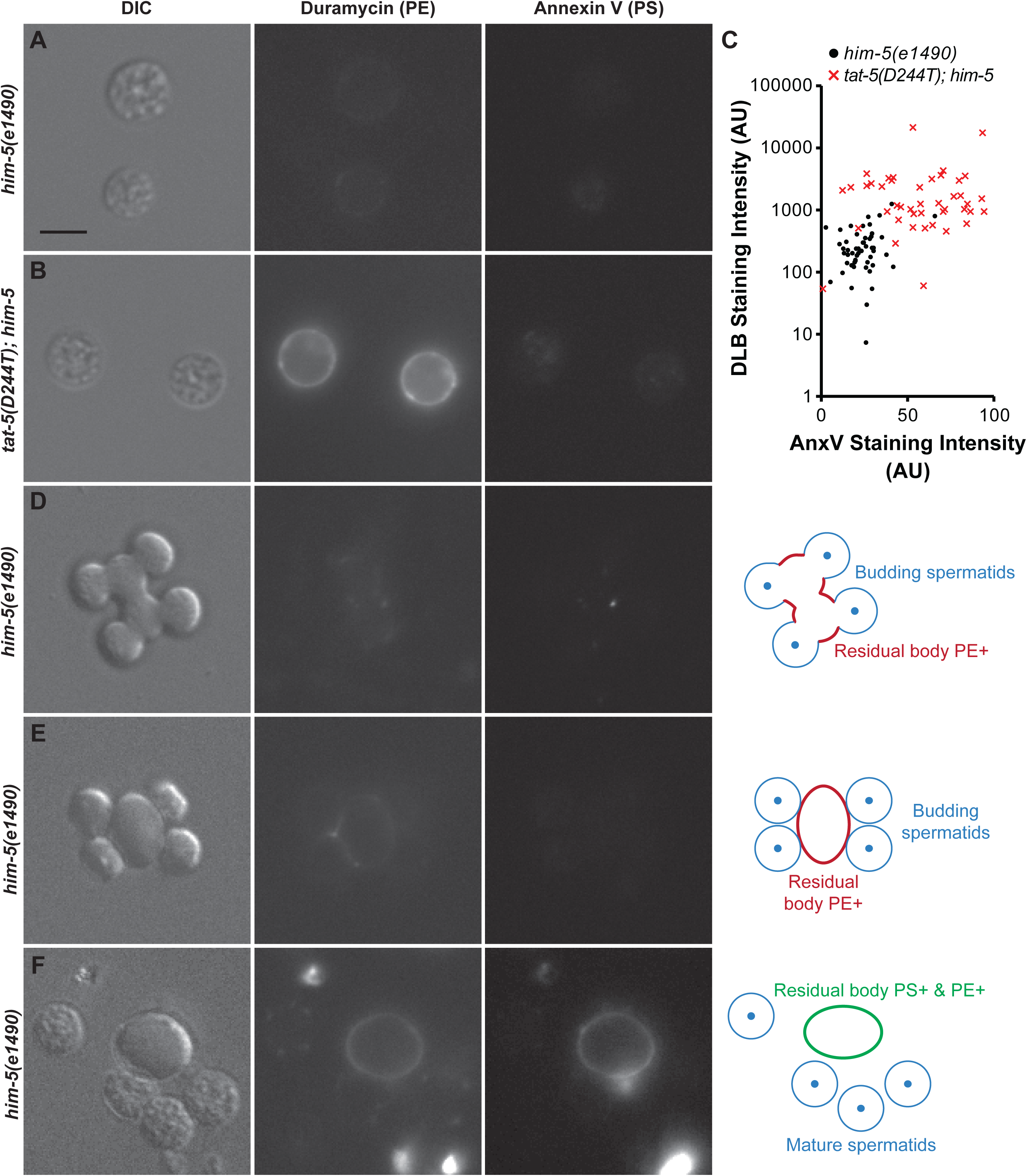
TAT-5 is required for phosphatidylethanolamine asymmetry in mature spermatids. A-B. Representative images of *him-5(e1490)* (A) or *tat-5(D244T); him-5(e1490)* (B) mature spermatids after duramycin and annexin V staining. Scale bar is 5 µm. C. Surface intensity of Annexin V and duramycin fluorescence over background. Each dot represents a single mature spermatid. D-F. Representative images of *him-5(e1490)* budding spermatids and residual bodies after duramycin and annexin V staining.

To confirm whether *tat-5* mutant spermatids specifically lose PE asymmetry, we co-stained dissected sperm with AnnexinV, which binds preferentially to exposed phosphatidylserine (PS) (Thiagarajan and Tait, 1990). We did not observe plasma membrane labeling on either *him-5* control or *tat-5(D244T); him-5* spermatids (Figure 4A-B). However, we could measure a two-fold increase in green fluorescence on the surface of *tat-5(D244T); him-5* spermatids (Figure 4C). We also did not observe a clear correlation between Duramycin and AnnexinV staining intensity (Figure 4C), confirming that PE exposure does not always correlate with PS exposure. Therefore, TAT-5 maintains PE asymmetry in the male germ line, not PS asymmetry.

### PE is exposed on residual bodies during meiosis

Given the localization of TAT-5 to the residual body membrane during meiosis (Figure 3E), we analyzed the timing of PE and PS exposure during residual body formation. We dissected *him-5* males, stained sperm with duramycin and Annexin V, and imaged developing spermatids in the process of budding from a residual body. We observed weak labeling of the forming residual body with duramycin (Figure 4D-E), which stained brighter as the spermatids separated from the residual body (Figure 4G, n=14). Similar to mature spermatids (Figure 4A), we only observed weak puncta of duramycin along the membrane of budding spermatids (Figure 4E). These data suggest that PE asymmetry is lost gradually and specifically on the residual body membrane. Furthermore, these asymmetric and symmetric lipid domains appear to be kept separate in the shared plasma membrane during spermatogenesis.

We next compared the timing of PE exposure to PS exposure. Consistent with PS-mediated phagocytic clearance of residual bodies (Huang et al., 2012), we did observe Annexin V labeling the residual body membrane after its release from mature spermatids (Figure 4F). However, in contrast to duramycin, both budding spermatids and forming residual bodies appeared unstained by Annexin V (Figure 4D-E). Together, these results indicate that the residual body membrane loses PE asymmetry before PS asymmetry.

During our staining experiments, we also observed bright puncta near the residual body labeling with both duramycin (n=10/14) and Annexin V (n=8/14) (Figure 4D, 4F), consistent with a stronger loss of lipid asymmetry on small extracellular vesicles released during spermatogenesis or dying cell fragments resulting from dissections. Thus, lipid asymmetry appears to be regulated differentially in budding spermatids, the residual body, and dying cells.

### TAT-5 is required for male fertility

Given the observed dynamics of TAT-5 localization and PE asymmetry, we next sought to determine the impact of TAT-5 and disrupted PE asymmetry on sperm maturation and function. To generate mutant males, we crossed the null and partial loss-of-function *tat-5* alleles to *him-5(e1490)* mutants. Examination of males after DAPI staining showed that both the *tat-5(tm1741)* null and *tat-5(D244T)* hypomorphic mutant germ lines appear normal, producing many sperm (brackets in Figure 5A, n=10). Thus, sperm appear to have progressed through meiosis to form spermatids in *tat-5* mutant males.

**Figure 5.**
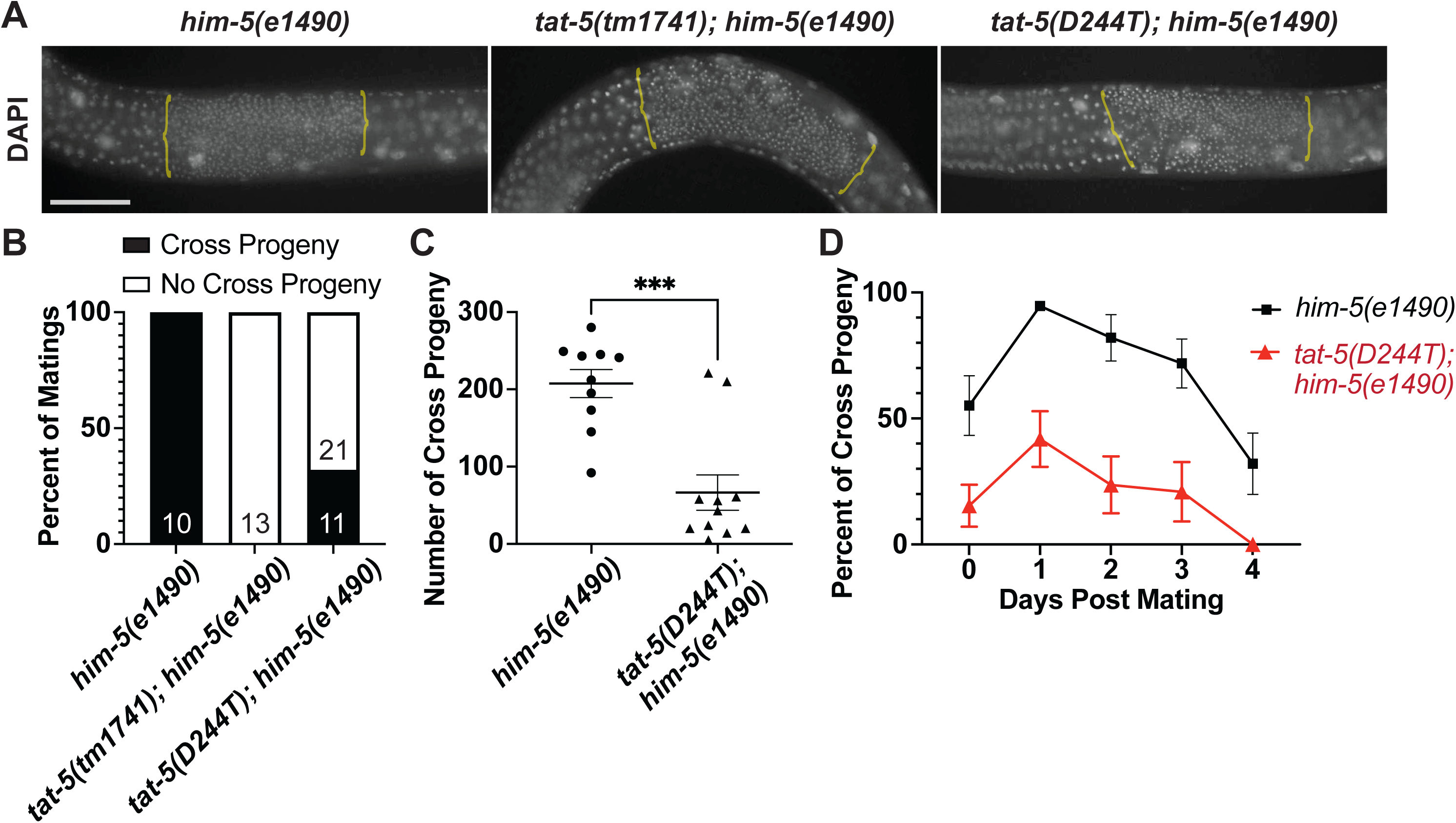
TAT-5 is required for male fertility. A. Representative images of DAPI staining of *him-5(e1490), tat-5(tm1741); him-5(e1490),* and *tat-5(D244T); him-5(e1490)* males. Brackets indicates region with mature spermatids. Scale bar is 20μm B. Percentage of matings with *dpy-11(e224)* hermaphrodites generating cross progeny, n=10, 12, and 32 crosses. C. Number of cross progeny after successful mating with *dpy-11(e224)* hermaphrodites for *him-5(e1490)* n=10 or *tat-5(D244T); him-5(e1490)* n=11 males. D. Percentage of cross progeny generated after mating males with *dpy-11(e224)* hermaphrodites. B. Fisher’s exact test, p<0.0001. C-D. Error bars ± SEM. One way ANOVA, *** p<0.001.

To understand how sperm contribute to the fertility phenotypes observed for hermaphrodites (Figure 2), we mated *tat-5; him-5* mutant males with a *dpy-11(e224)* hermaphrodite and observed their ability to sire non-Dumpy progeny. Control *him-5* males generated cross progeny 10 out of 10 times (Figure 5B). In contrast, we found that *tat-5(tm1741)* null mutant males were unable to generate cross progeny (n=11) (Figure 5B). Hypomorphic *tat-5(D244T)* males also failed to sire cross progeny two-thirds of the time (n=21/32), siring significantly less frequently than *him-5* males. These defects could be due to decreased function of *tat-5* mutant sperm or failure of *tat-5* mutant males to mate.

As *tat-5(D244T)* males sire cross progeny one-third of the time (n=11/32), we could analyze the fitness of sperm after a successful mating with *dpy-11(e224)* hermaphrodites. Matings with hypomorphic *tat-5(D244T)* males produced significantly fewer cross progeny compared to *him-5* control matings (Figure 5C), with most matings producing only one-eighth the cross progeny (n=9/11). When the mating data was analyzed daily, we found that sperm from wild type males primarily produced cross progeny for the first few days after mating, with self-progeny first dominating 4 days after mating (Figure 5C). In contrast, the *tat-5* hypomorph consistently produced less than 50% cross progeny every day after a successful mating (Figure 5D). These data suggest that male sperm with reduced TAT-5 activity were not able to compete with hermaphrodite sperm.

### TAT-5 is required for sperm activation and migration

For male sperm to outcompete hermaphrodite sperm after mating, male sperm first migrate inside the hermaphrodite gonad from the vulva to the spermatheca. To assess sperm migration, males were mated with feminized strains unable to produce sperm. When raised at the restrictive temperature, *fem-1(hc17ts)* mutants produce no sperm, allowing us to use DAPI staining to assess the number of mated *fem-1(hc17ts)* mutants that had sperm transferred and localized to the spermatheca (Figure 6A). As expected, unmated *fem-1* mutants showed no sperm in the spermatheca at the restrictive temperature (n=10, Figure 6B, 6F), while mating with *him-5(e1490)* resulted in many sperm in the spermatheca in 93% of matings (n=14/15, Figure 6C, 6F). In contrast, after overnight mating with *tat-5* mutant males, we never observed many sperm in the *fem-1* spermatheca (Figure 6F). Two-thirds of *tat-5* null mutant (n=7/11) or *tat-5(D244T)* mutant (n=7/10) matings resulted in no sperm detected in the spermatheca by DAPI staining (Figure 6D, 6F). However, one-third of *tat-5* null mutant (n=3/11) or *tat-5(D244T)* mutant (n=3/10) matings without visible sperm did result in fertilized embryos (Figure 6F), which indicates that at least a few sperm were transferred. In cases where both sperm and embryos were present (Figure 6E-F), we sometimes observed sperm in the *fem-1* uterus after mating with *tat-5* mutant males. These results suggest that *tat-5* mutant sperm rarely take up residence in the spermatheca, even in the absence of competing hermaphrodite sperm.

**Figure 6.**
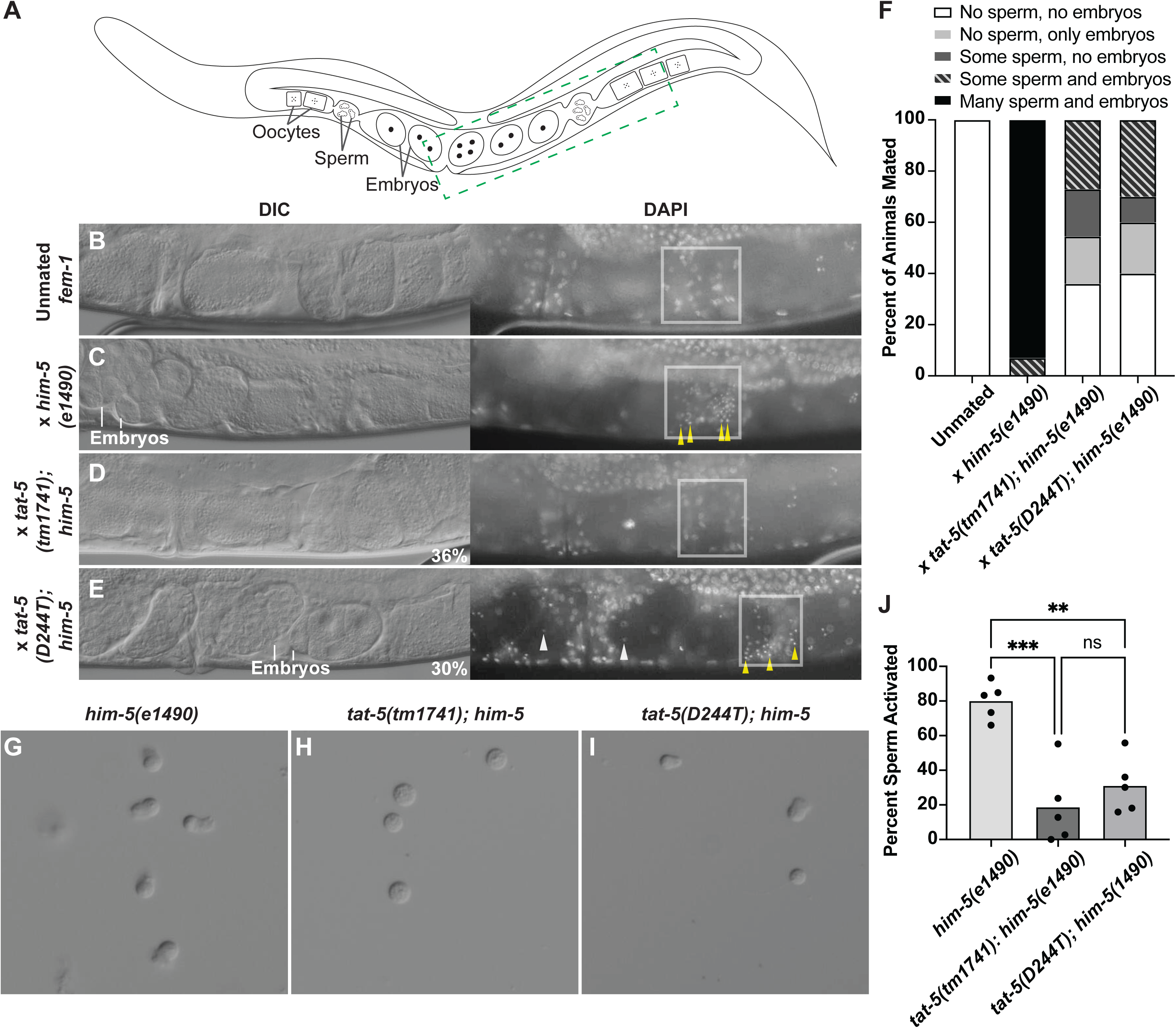
TAT-5 is required for spermiogenesis. A-E. Schematic (A) and representative images of DAPI sperm transfer to *fem-1(hc17ts)* hermaphrodite animals (B) after mating with *him-5(e1490)* (C)*, tat-5(tm1741); him-5(e1490)* (D), or *tat-5(D244T); him-5(e1490)* (E) males. A few representative spermatids in the spermatheca (yellow arrowheads) and uterus (white arrowheads) are indicated. F. Prevalence of sperm transferred to *fem-1(hc17ts)* hermaphrodites. n≥10 for all groups. G-I. Representative images of sperm after activation with Pronase treatment. J. Quantification of sperm activation based on morphology. Each dot represents the sperm resulting from two dissected males (n=10 males dissected for each genotype).Statistical significance was calculated using the Mann-Whitney U test.

To understand why *tat-5* sperm were unable to reach or stay in the spermatheca, we examined the morphology of mature spermatids dissected in sperm media. Although dissected *tat-5* mutant spermatids appeared mostly normal, we noticed that there were changes to nuclear positioning in 10% of *tat-5* mutant spermatids, a six-fold increase in off-center nuclei over *him-5* controls (Table 1). Off-center nuclei were previously observed in one-third of *spe-10* mutant spermatids (Shakes and Ward, 1989b), which is intriguing because *spe-10* mutant sperm are defective in forming a migratory pseudopod. However, whether the off-center nuclei phenotype contributes to sperm migration defects is unclear.

**Table 1.**
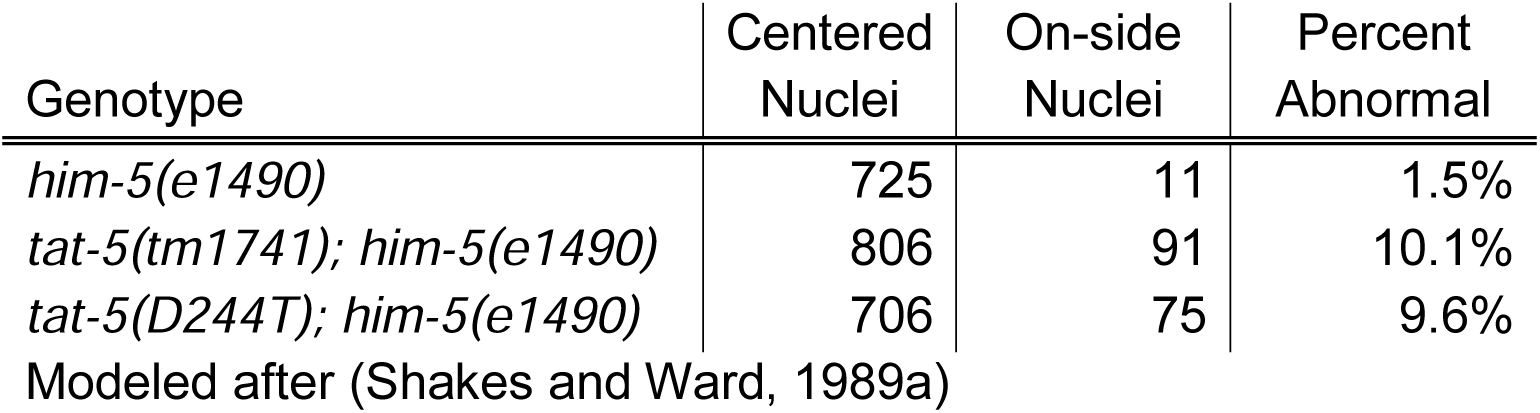
Nuclear Position in Spermatids.

For sperm to migrate, they need to activate and transition from a round immotile spermatid to a motile spermatozoon with a pseudopod (Shakes and Ward, 1989a). Therefore, we wanted to know if *tat-5* mutant sperm were able to change their morphology in the presence of an *in vitro* sperm activator, Pronase. While 80% of dissected control *him-5* sperm activated and formed pseudopods after Pronase treatment (Figure 6G, 6J), only 20-30% of *tat-5* mutant sperm formed a pseudopod (Figure 6H-J). Most *tat-5* mutant sperm maintained a round immotile morphology in the presence of Pronase (Figure 6H), suggesting that *tat-5* mutant sperm have defects in receiving the activation signal or changing their cell shape to form a pseudopod. Difficulty activating or migrating can explain why *tat-5* mutant sperm are unable to establish or maintain their residence in the spermatheca after mating.

## Discussion

This work describes the first role for a P4B-ATPase in sperm and confirms the importance of TAT-5 for oogenesis. We demonstrated that a loss of *tat-5* decreases male and hermaphrodite fertility by disrupting both oocyte production and sperm activation, although only oogenesis can proceed with a low level of TAT-5 activity. Our results suggest that TAT-5 flippase activity is critical during spermatogenesis to establish PE asymmetry on the spermatid plasma membrane, in addition to the female germ line (Beer et al., 2018; Wehman et al., 2011). We demonstrated that even with low levels of TAT-5 activity, PE was exposed on the surface of mature spermatids that were unable to activate to form a motile pseudopod (Figure 7). Our work, combined with others, demonstrates the importance of regulating membrane asymmetry in sperm (Flesch and Gadella, 2000).

**Figure 7.**
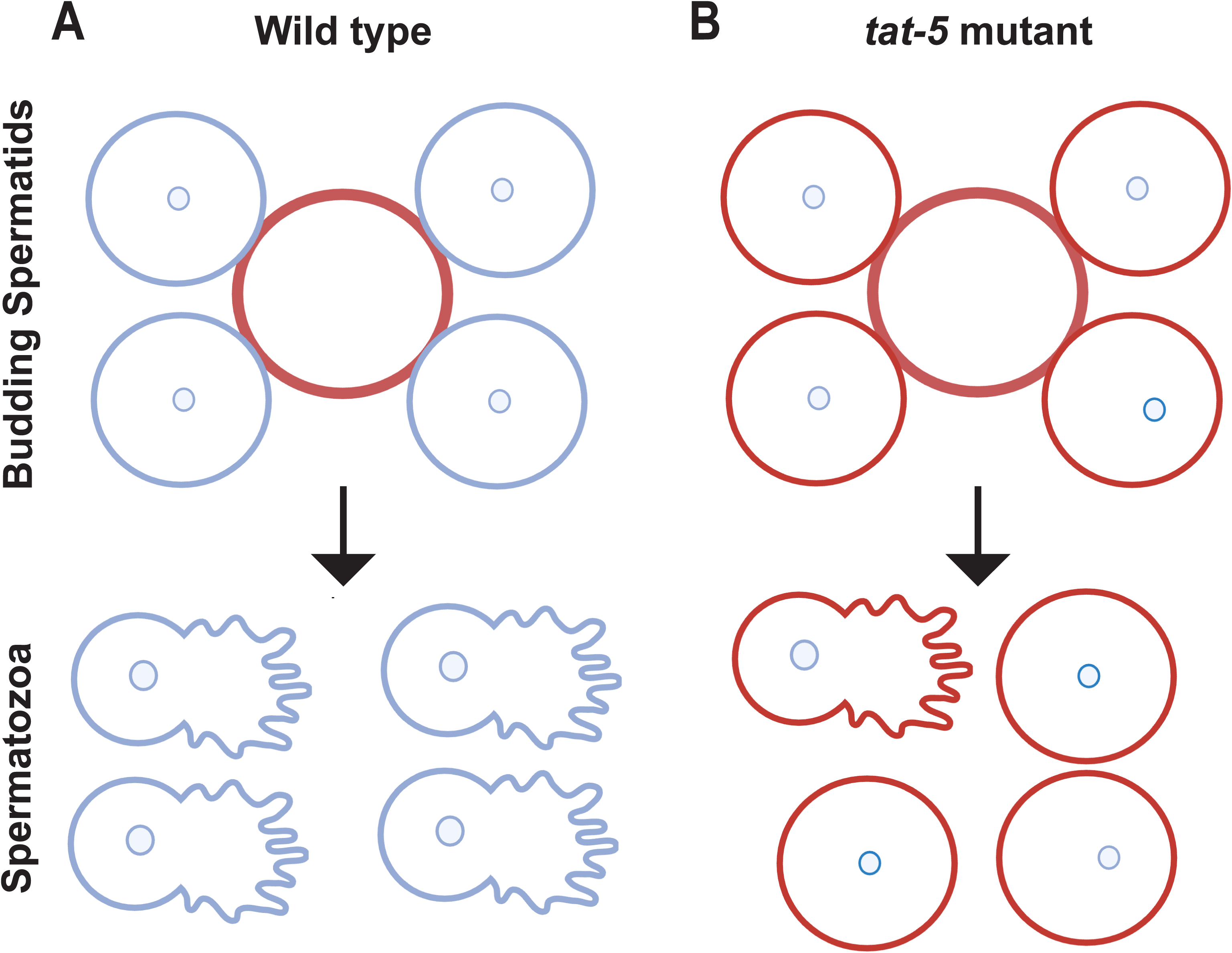
Model for the role of TAT-5 in sperm. A. TAT-5 internalizes phosphatidylethanolamine (PE) on the surface of primary and secondary spermatocytes, establishing lipid asymmetry for the spermatid plasma membrane (blue outline) and allowing normal activation and migration. PE is normally exposed on the residual body membrane (red outline). B. Loss of TAT-5 activity results in PE exposure on the outside of the spermatid membrane (red outline), off-center nuclei, and defects in sperm activation to form pseudopods.

Mammalian P4A-ATPases also regulate lipid asymmetry in sperm and play a role in male fertility. Both ATP8B3 and ATP8B5 show testis-specific expression and localize to the acrosomal region of sperm (Gong et al., 2009; Wang et al., 2004; Xu et al., 2009). The acrosome is a vesicle in the sperm head that fuses with the plasma membrane to release proteins that help the sperm penetrate the zona pellucida and reach the egg (Bianchi and Wright, 2020). ATP8B3 mutant sperm fail to bind to or penetrate the egg and prematurely expose PS on their surface (Wang et al., 2004), while ATP8B5 is thought to flip PC and PE lipids (Xu et al., 2009), suggesting a potential role for lipid asymmetry in the acrosome reaction. Given the infertility of ATP9A knockout mice (Meng et al., 2023a), it will be important to determine how ATP9A and ATP9B, the mammalian P4B-ATPases homologous to TAT-5, promote male fertility.

Cytosolic proteins that are not needed during sperm activation or fertilization are typically segregated into the residual body (Nelson and Ward, 1980), suggesting that the multipass transmembrane protein TAT-5 is intentionally trafficked away from maturing spermatids to end up in the residual body membrane. TAT-5 is the first transmembrane protein known to be specifically localized to the membrane of the forming residual body, suggesting that not only cytosolic components are segregated to the residual body during meiosis. Furthermore, our lipid staining experiments demonstrated that lipid asymmetry is specifically and progressively lost in the residual body and not in budding spermatids, suggesting the existence of lateral sorting mechanisms for lipids and proteins in the membrane during spermatogenesis. Therefore, GFP::TAT-5b will be a useful marker to investigate the mechanisms of membrane sorting during male meiosis. Furthermore, TAT-5 may help localize other proteins or organelles to the residual body, based on the known roles of TAT-5 homologs in endolysosomal trafficking (Hua et al., 2002; Hua and Graham, 2003; McGough et al., 2018; Meng et al., 2023b; Tanaka et al., 2016; Wicky et al., 2004).

Given that GFP::TAT-5b appeared to label hermaphrodite sperm in the spermatheca of intact hermaphrodites, we were surprised that GFP::TAT-5b appeared absent from mature spermatids in intact or dissected males. However, removal of residual bodies is more efficient in males compared to hermaphrodites and residual bodies have been observed in the spermatheca (Huang et al., 2012), raising the possibility that the GFP::TAT-5b we observed in the hermaphrodite spermatheca may be due to TAT-5 localization to residual bodies or the spermatheca itself.

We were also surprised to discover that residual bodies began losing PE asymmetry during their formation, despite the localization of the PE flippase TAT-5 to residual bodies. This suggests that TAT-5 activity establishes the asymmetry of the spermatid membrane during the primary and secondary spermatocyte stages, but that TAT-5 is quickly inactivated in residual bodies during spermatogenesis. As mitochondria are sorted into the budding spermatids and away from the residual body, this could lead to regional differences in ATP availability (Huang et al., 2012). In addition, the rapid PE exposure suggests that lipid scramblases are quickly activated in forming residual bodies to disrupt lipid asymmetry. Notably, PE exposure preceded PS exposure, suggesting different regulation of PS flippases, PE flippases like TAT-5, and non-specific lipid scramblases. Determining how flippase and scramblase activities are controlled to disrupt lipid asymmetry in residual bodies will be important to understand how cells balance the structural and signaling roles of lipid asymmetry.

Although we discovered a fine choreography of lipid localizations during spermatogenesis, the role that PE asymmetry plays during sperm activation is unclear. One possible explanation for the loss of pseudopod formation in *tat-5* mutant sperm is an inability to sense the activation signal with PE exposed, which could be due to the mislocalization of signaling proteins. However, given the unusually rounded morphology of cells in *tat-5* mutant embryos (Wehman et al., 2011), the round morphology of mutant sperm could also be due to cytoskeletal defects or lipid-induced changes to the biophysical properties of the membrane itself. Therefore, it will be important to determine how lipid asymmetry regulates sperm for fertilization, as these lipid functions are likely to be conserved in other developmental and pathogenic cell fusion events.

## Acknowledgements

We thank the Waksman Institute Shared Imaging Facility, Rutgers, and The State University of New Jersey for microscopy service particularly Nanci Kane for her help. Some strains were provided by the CGC, which is funded by NIH Office of Research Infrastructure Programs (P40 OD010440).

## Funding Sources

Research reported in this publication was supported by the National Institute of General Medical Sciences of the National Institutes of Health under award number R35GM152234 to A.M.W. and an NIH Institutional Research and Academic Career Development award (K12GM093854) fellowship to K.A.M. The National Institute of Child Health and Human Development also supported this research under award number R01HD054681 to A.W.S. and a K99 Pathway to Independence Award (K99HD115785) to K.A.M.

## Figure Legends

**Supplemental Figure 1.** Mature spermatids from control *him-5(e1490)* males imaged with conditions identical to Figure 3F. Scale bar represents 5 μm.

**Supplemental Table 1.**
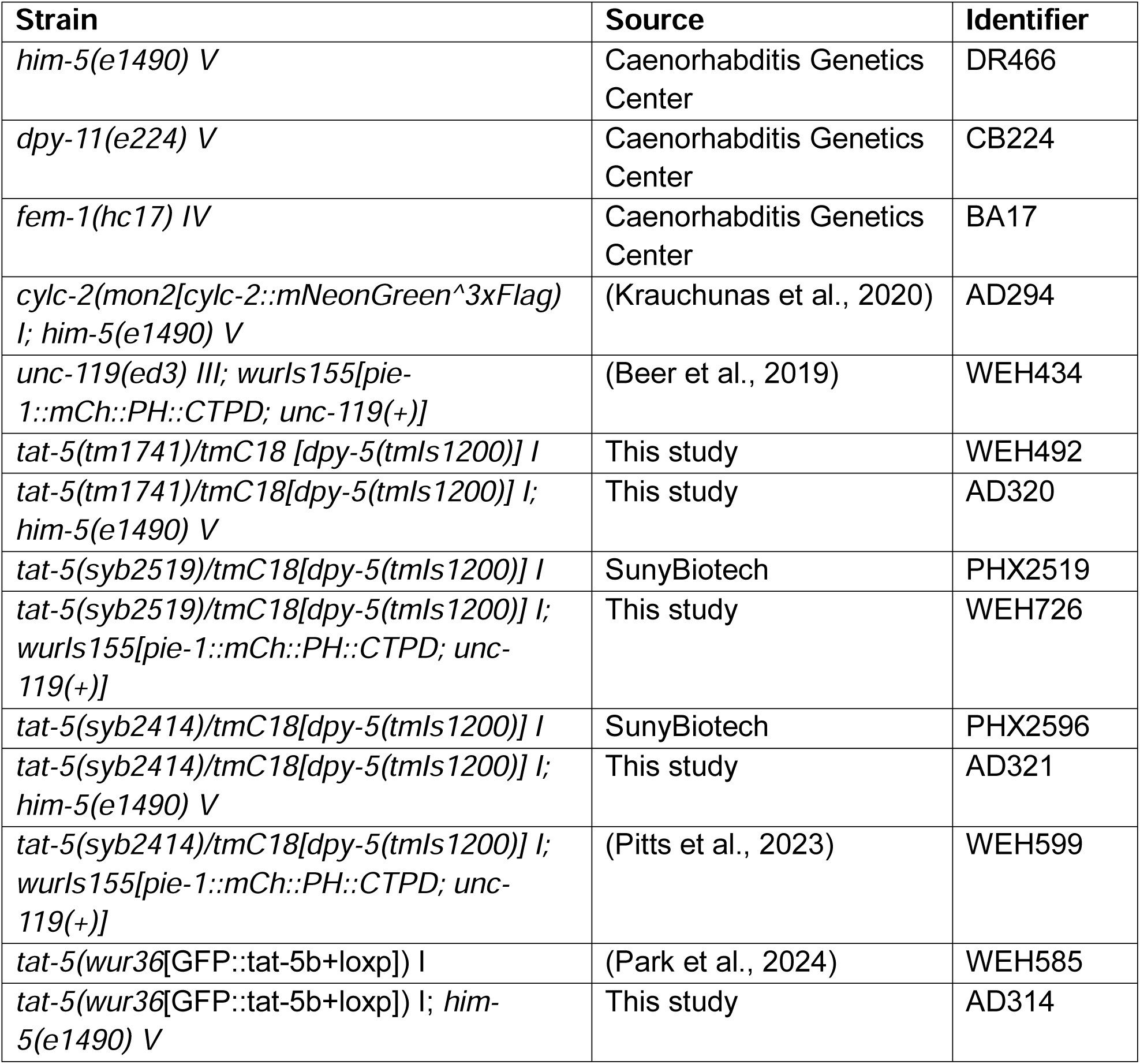
Strain List.

**Supplemental Table 2.**
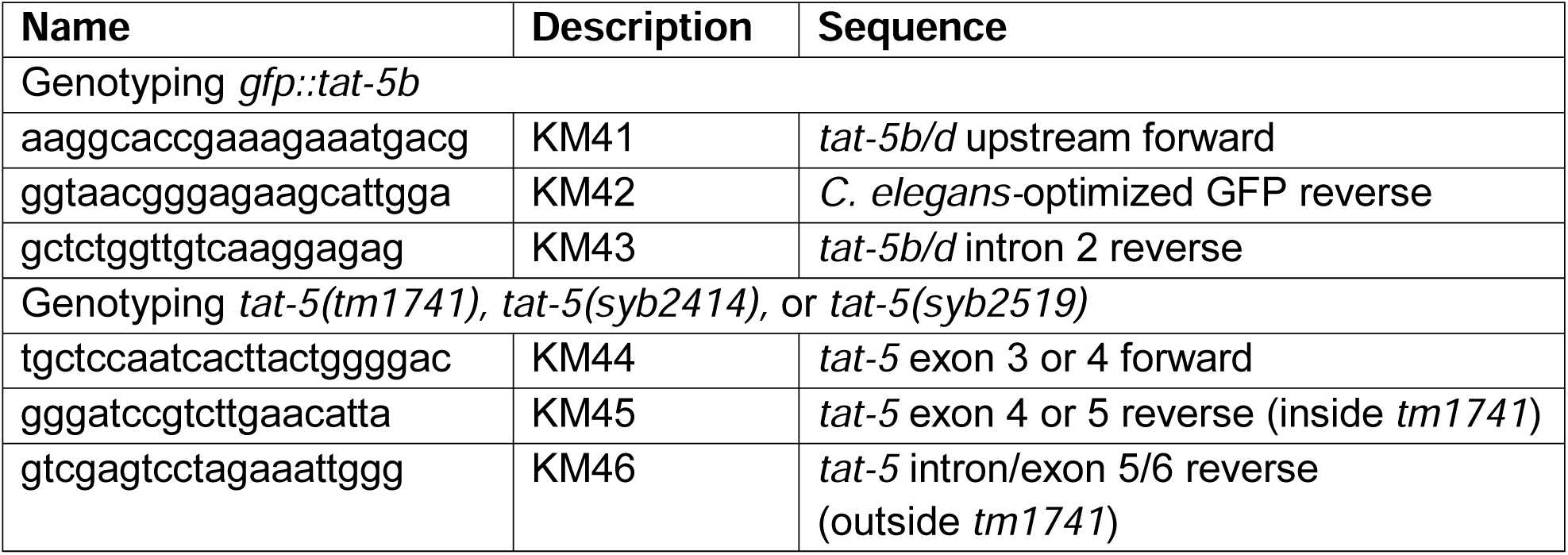
Primer List.

## Notes

### Competing Interest Statement

The authors have declared no competing interest.

