## Supplementary figures and images for "Sperm activation for fertilization requires robust activity of the TAT-5 lipid flippase"

### Supplemental Figure 1

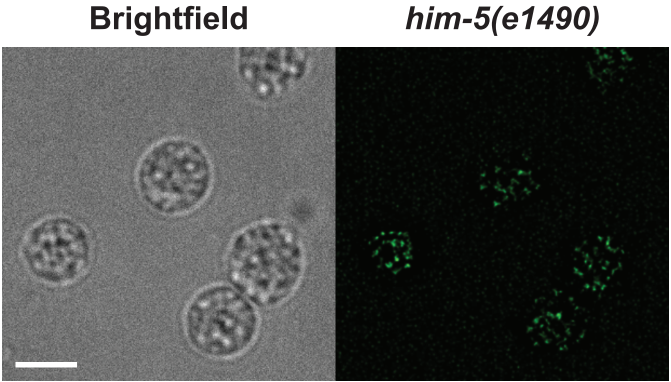
